# The Effect of Ex-vivo Gravid Uterine Support on Umbilical Vessel Flow Velocities and Ductus Arteriosus Patency

**DOI:** 10.1101/2025.03.14.643369

**Authors:** Jennifer M. Schuh, Araceli A. Morelos, Emmanuel L. Abebrese, Deron W. Jones, Natalie Stocke, Nicholas Starkey, Jose H. Salazar

**Author notes:** Respective email addresses. Contact Information: Jose H. Salazar, MD, PhD, Division of Pediatric Surgery, Medical College of Wisconsin, Children’s Corporate Center, Suite C320, 999 N 92nd St, Milwaukee, WI 53212.

## Abstract

**Purpose:** Fetal lamb artificial womb (AW) models rely on perfusion delivered via extracorporeal membrane oxygenation (ECMO) through umbilical or central vessel cannulation. Ductus arteriosus (DA) patency is paramount to fetal survival but may be disrupted by physiological stress and ECMO support. Existing AW models without the placenta use prostaglandin infusion to maintain DA patency, risking side effects like dysrhythmias, edema, or hypotension. We aim to investigate DA patency in an ex-vivo sheep model of gravid uterine support and quantify placental regulation of umbilical vessel flow with extrauterine flow variations.

**Methods:** An ex-vivo placental perfusion model was employed with bilateral uterine artery and vein cannulation. To maintain adequate flow, there is frequent manual titration of pump revolutions per minute, altering the blood pressure delivered to the uterus. Ultrasound measurements were obtained before laparotomy and on ECMO to document shunt patency and umbilical vessel and DA flow velocities; velocities were compared with student’s paired t-tests.

**Results:** Six of eleven sheep had ultrasounds obtained on full ECMO support. There was no difference in peak average velocity through the ductus before or after ECMO cannulation (*p*=0.26; CI -0.41 – 0.13), umbilical vein peak velocity (*p*=0.78; CI -0.08 – 0.10), or umbilical artery peak velocity (*p*=0.73; CI -0.17 – 0.13).

**Conclusion:** While perfusing a gravid uterus via the uterine vessels, we demonstrate a patent fetal DA and relatively constant umbilical vein flow velocities despite fluctuations in the ECMO pressure delivery to the uterus. The placenta therefore maintains regulation of umbilical vein blood flow velocity, facilitating physiologic shunt patency.

## INTRODUCTION

Prematurity is the leading cause of neonatal mortality. For every ten live births in the United States, one is premature (defined as <37 weeks of gestational age) [1]. The risk of mortality of a premature newborn drops significantly with each week that the gestation approximates term, from >80% mortality for an infant born at 22 weeks of gestation to <10% for one born at 28 weeks [2], [3], [4]. Survival is not the only relevant outcome, as most of the periviable infants that live to be discharged have neurodevelopmental impairment, a risk that also decreases with longer gestational age (GA) [4]. One approach to improving periviable neonatal outcomes is establishing a system that allows for the continuation of an “in-utero-like” environment, enabling the conservation of fetal circulation. This idea has existed for decades [5], [6], [7]. Fetal lamb artificial womb (AW) and artificial placenta (AP) models rely on perfusion delivered via extracorporeal membrane support (ECMO) through umbilical or central vessel cannulation in the absence of the placenta. Consistent, prolonged survival has been achieved for up to four weeks with the advent of low-resistance oxygenators that permit the fetal heart to send blood through a circuit without reliance on a pump [8], [9], [10], [11].

The patency of fetal circulation is paramount to prolong the fetus’s survival prior to exposure to an air environment. Physiological stress and ECMO support may disrupt the physiologic environment required to maintain ductus arteriosus (DA) patency. Specifically, the cardiac preload is altered by increased pressures delivered by the ECMO pump, to a lesser degree with pumpless models, and afterload is increased in pumpless models relying on the fetal heart to distribute blood through the circuit tubing. Existing AW and AP models use continuous infusion of prostaglandins to maintain DA patency, exposing the fetus to side effects like dysrhythmias, edema, or hypotension [10], [12], [13]. The model of an artificial womb with placental preservation, first described in 2014, may temper these limitations by maintaining the placental barrier between the circuit and fetal circulation [14]. It is unknown whether placental preservation is sufficient to maintain DA patency in the absence of prostaglandins. Further, it is unknown to what extent the placenta in this model regulates blood flow to the umbilical vessels.

Our primary aim was to investigate fetal DA patency in an ex-vivo sheep model of gravid uterine support. Secondarily, we aimed to quantify placental regulation of umbilical vessel flow with extrauterine flow variations. We hypothesized that placental preservation would permit DA patency without the use of prostaglandins, and that umbilical vessel blood flow would be consistently maintained by the placenta despite extrauterine flow variations.

## METHODS

### Data source

Eleven gravid ewes were used, of which six had ultrasounds obtained on full ECMO support. All animals were maintained under controlled temperatures with access to food and water. The ewes were fasted overnight prior to anesthesia. Fetuses were between 110 and 115 days old, corresponding to periviability [14].

### Experiment

An ex-vivo placental perfusion model was employed; protocol was based on Salazar et al. [14]. The ewe was induced with general anesthesia, 200mg rectal indomethacin was given as a tocolytic, and a low transverse laparotomy was performed. The uterus was delivered, and bilateral uterine arteries and veins were isolated and cannulated. Time on full ECMO support was recorded when all uterine vessel flow was supplied by the ECMO circuit. Transverse stapled cervical hysterectomy was completed, and the uterus was transferred to a warmed tank of lactated ringers. To maintain adequate flow, there was constant manual titration of pump revolutions per minute, leading to fluctuations in the pressure by which blood is delivered to the uterus. Prostaglandins were not administered. Doppler ultrasound measurements were obtained before laparotomy and at least two hours after transition to ECMO support to document shunt patency and flow velocities at the umbilical cord vessels and DA. The circuit was primed with whole blood obtained from different donor ewes, and blood or crystalloid was administered for volume maintenance. Electrolytes, glucose, and activated clotting time were monitored and corrected as needed. If the fetus survived for >2 additional hours, ultrasound measurements were repeated.

### Outcomes

#### Primary

The primary outcome was difference in DA peak velocity between measurements at baseline and measurements on ECMO circuit.

#### Secondary

The secondary outcome was difference in umbilical vessel peak velocity between measurements at baseline and measurements on ECMO circuit.

### Statistical analysis

Baseline velocities were compared to velocities on ECMO support with student’s paired t-tests. All analyses were performed using Microsoft Excel© (Redmond, WA).

### Ethics

All procedures were performed in accordance with relevant laws and institutional guidelines and have been approved by the Institutional Animal Care and Use Committee (AUA00007678, initial approval 12/8/2021).

## RESULTS

### Data source

Six gravid uteri with singleton pregnancies (mean 113 ± 1.3 days gestation) had ultrasounds obtained on full ECMO support. Mean time to ultrasound after full ECMO support was 3 hours, 9 minutes ± 1 hour, 8 minutes. Two gravid uteri had repeat ultrasounds at 4 hours, 15 minutes and 6 hours, 58 minutes after full ECMO support.

### Outcomes

There was no difference in peak average velocity through the DA before or after ECMO cannulation (*p*=0.26; CI -0.41 – 0.13), umbilical vein peak velocity (*p*=0.78; CI -0.08 – 0.10), or umbilical artery peak velocity (*p*=0.73; CI -0.17 – 0.13). Velocity differences from baseline are specified in Table 1. Representative images are shown in Figure 1.

**Table 1.** Ductus arteriosus and umbilical vessel velocity differences from baseline after ECMO support with placental preservation.

| Sheep | Sex | Gestational age (days) | Time on ECMO (hh:mm) | Ductus arteriosus peak average velocity difference (m/s) | Umbilical vein peak velocity difference (m/s) | Umbilical artery peak velocity difference (m/s) |
| --- | --- | --- | --- | --- | --- | --- |
| 1 | M | 113 | 2:51 | 0.47 | 0.30 | 0.40 |
| 2 | F | 113 | 2:02 | 0.54 | 0.10 | -0.08 |
|  |  |  | 4:15 | 0.08 | -0.01 | -0.23 |
| 3 | M | 114 | 3:35 | -0.06 | 0.14 | 0.37 |
|  |  |  | 6:58 | 0.46 | 0.00 | 0.23 |
| 4 | F | 110 | 2:24 | -0.29 | -0.16 | 0.01 |
| 5 | F | 113 | 2:51 | 0.15 | -0.18 | -0.18 |
| 6 | M | 114 | 5:14 | 0.22 | 0.01 | 0.06 |

Abbreviations: ECMO- extracorporeal membrane oxygenation; hh-hours; mm- minutes; s-seconds; m- meters

**Figure 1.**
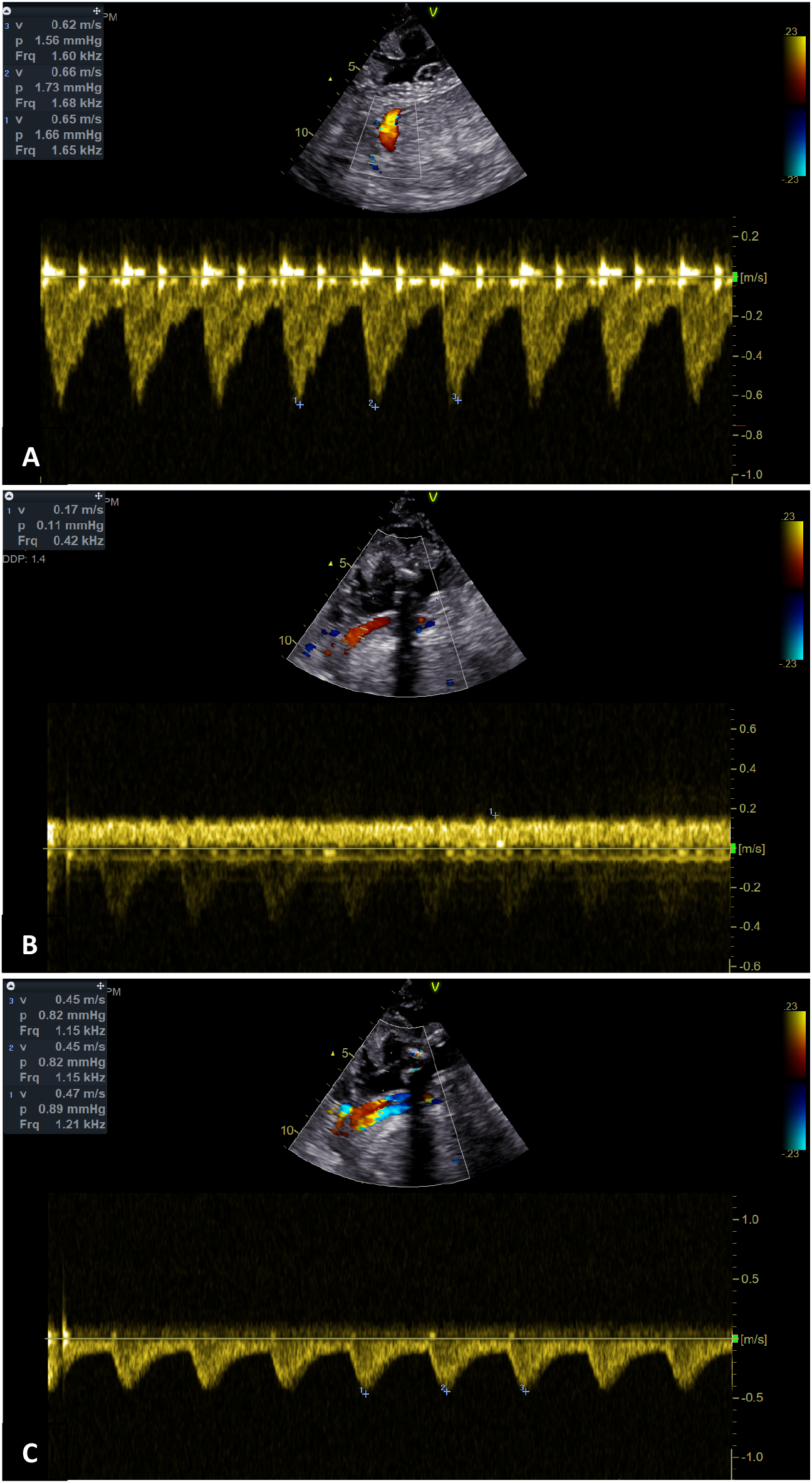
Representative images of fetal ultrasounds after five hours on ECMO (A) Ductus arteriosus (B) Umbilical vein (C) Umbilical artery Abbreviation: ECMO-extracorporeal membrane oxygenation

## DISCUSSION

In this placental preservation AW model, ductus arteriosus patency was reproducibly established via ultrasound after transition to full ECMO support. There was no significant difference in blood flow velocity from baseline (before laparotomy) to >2 hours after full ECMO support within the ductus arteriosus, umbilical artery or umbilical vein, despite alterations in the blood pressure being delivered from the ECMO circuit to the uterine vessels. Therefore, the placenta plays a key role in facilitating shunt patency and regulating blood flow to the fetus.

Uteroplacental circulation has previously been identified as a key component of healthy pregnancies in humans, but relatively little is known about the placental regulation of blood flow to the fetus [15], [16]. Uteroplacental perfusion is normally facilitated by low placental resistance and increased maternal cardiac output [17]. It has been established in a gravid ovine model that ewes with increased uterine artery blood flow throughout gestation (facilitated by resveratrol administration) have increased fetal weight and absolute fetal oxygen delivery, without altering fetal blood pressure or hemodynamics, including blood pressure and velocity measurements in the ductus arteriosus and umbilical vein, compared to a control group with lower uterine artery blood flow [18], [19]. Our findings are consistent in demonstrating preserved blood flow and velocity to the umbilical vessels and ductus arteriosus with uterine artery flow variability. We add the advantage of measuring pre- and post-artery flow variability in the same uteri, providing a direct comparison of continuous titration that speaks directly to the physiological function of the placenta relative to a single pregnancy. Furthermore, we directly adjust the blood pressure delivery to the gravid uterus and measure resultant fetal flows.

The placental regulation of blood flow to the fetus has substantial repercussions in the artificial womb model. Fetal circulation patency is quintessential to facilitate fetal survival prior to the transition to oxygenation via pulmonary air exchange but is threatened by the biological stress of ECMO. The prostaglandin infusions currently utilized by AW models risk substantial arrhythmogenic and hemodynamic fetal side effects [10], [12], [13]. Therefore, demonstrating the ability of the placenta to maintain a patent DA, with no significant difference in flow velocity in vivo compared to on full ECMO support, offers a significant advantage of placental preservation.

This study has considerable implications but must be interpreted in the context of select limitations. Due to the resource intensive nature of large animal experimentation, the sample size is relatively small. However, even so, pre- and post-velocities for each parameter measured resulted in a significant confidence interval, suggesting adequacy of the sample. The sample may be biased by the measurements we were unable to obtain; that is, the five uteri which were not able to be measured >2 hours after transition to full ECMO support may have altered the findings, though we feel this is unlikely. Further, though the patency of the DA and umbilical flow velocities speak to the placental regulation of fetal perfusion, they are not necessarily perfect correlates for nutrient delivery, which ultrasound is unable to measure.

## CONCLUSION

We demonstrate patent ductus arteriosus and relatively constant umbilical vessel flow velocities in an ex-vivo placental perfusion artificial womb model, despite fluctuations in the circuit blood flow and pressure delivery to the uterus. The placenta therefore maintains regulation of umbilical vein blood flow velocity, facilitating physiologic shunt patency.

## GLOSSARY

- Extracorporeal membrane oxygenation- a support modality in which blood is mechanically oxygenated in a circuit before returning to the individual/tissue
- Ductus arteriosus- a fetal blood vessel connecting the pulmonary artery to the aorta, physiologically closes at birth

## ACKNOWLEDGEMENTS

We acknowledge Lindsey Kalvin and the Medical College of Wisconsin Echocardiography Core for indispensable contributions acquiring images.

## AUTHOR CONTRIBUTIONS

Jennifer M. Schuh-conceptualization, data curation, formal analysis, investigation, methodology, visualization, writing-original draft, writing-reviewing and editing Araceli A. Morelos-conceptualization, data curation, formal analysis, methodology, visualization, writing-reviewing and editing Emmanuel L. Abebrese-conceptualization, data curation, formal analysis, methodology, visualization

Deron W. Jones-data curation, methodology, visualization, writing-reviewing and editing Natalie Stocke-conceptualization, data curation, formal analysis, methodology, visualization, writing-reviewing and editing Nicholas Starke - conceptualization, data curation, formal analysis, methodology, visualization, writing-reviewing and editing

Jose H. Salazar-conceptualization, data curation, formal analysis, investigation, methodology, project administration, supervision, validation, visualization, writing-reviewing and editing All authors approve of the final manuscript and agree to be accountable for all aspects of the work.

